# Estimating the Probability of Early Afterdepolarization and Predicting Arrhythmic Risk associated with Long QT Syndrome Type 1 Mutations

**DOI:** 10.1101/2020.04.09.034843

**Authors:** Qingchu Jin, Joseph L. Greenstein, Raimond L. Winslow

## Abstract

Early after-depolarizations (EADs) are action potential (AP) repolarization abnormalities that can trigger lethal arrhythmias. Simulations using biophysically-detailed cardiac myocyte models can reveal how model parameters influence the probability of these cellular arrhythmias, however such analyses can pose a huge computational burden. We have previously developed a highly simplified approach in which logistic regression models (LRMs) map parameters of complex cell models to the probability of ectopic beats (EBs). Here, we extend this approach to predict the probability of early after-depolarizations (P(EAD)). We use the LRM to investigate how changes in parameters of the slow-activating delayed rectifier current (I_Ks_) affect P(EAD) for 17 different Long QT syndrome type 1 (LQTS1) mutations. We compare P(EAD) for these 17 LQTS1 mutations with two other recently proposed model-based arrhythmia risk metrics. These three model-based risk metrics yield similar prediction performance; however, they all fail to predict relative clinical risk for a significant number of the 17 studied LQTS1 mutations. The consistent successes and failures of all three risk metrics suggest that important functional characteristics of LQTS1 mutations may not yet be fully known.

**Author summary:** An early after-depolarization (EAD) is an abnormal cellular electrical event which can trigger dangerous arrhythmias in the heart. We use our previously developed method to build a simple logistic regression model (LRM) that estimates the probability of EAD (P(EAD)) as a function of myocyte model parameters. Using this LRM along with two other recently published model-based arrhythmia risk predictors, we estimate risk of arrhythmia for 17 Long QT syndrome type 1 (LQTS1) mutations. Results show that all approaches have similar prediction performance in that there are a set of mutations whose relative clinical risk for arrhythmia are well estimated using these metrics, but that relative risk is consistently over- or under-estimated across all approaches for a significant number of other mutations. We believe this indicates that the functional characterization of the LQTS1 phenotype is incomplete.

## Introduction

Early after-depolarizations (EADs) are abnormal repolarization events that occur during the plateau or rapid repolarization phase of action potentials (APs), which can in turn trigger large scale cardiac arrhythmias in the setting of long QT syndrome [1–3]. Computational modeling of EADs has provided insights into the mechanisms by which they can trigger arrhythmias in the heart [4–6].

We seek to understand how variations in the underlying biophysical properties or physiological state of the myocyte influence the probability of occurrence of EADs. Stochastic mechanisms play a role in EAD initiation. Fowler et al. demonstrated that stochastic Ca^2+^ sparks and local Ca^2+^ waves can generate sufficient Na^+^/Ca^2+^ exchanger current to trigger EADs [7]. Tanskanen et al. also showed that a phosphorylation-induced change in LCC stochastic gating contributes to EAD initiation [8]. Therefore, we use a three-dimensional spatial model of the ventricular myocyte developed by Walker et al. [9] in which the fundamental events governing intracellular calcium (Ca^2+^) dynamics are modeled stochastically.

The complexity of this stochastic model makes it challenging to perform the many repeated simulations needed to estimate the probability of EADs, denoted P(EAD), as a function of underlying model parameters. Here, we develop a logistic regression model (LRM) that directly maps myocyte model parameters to P(EAD) [10]. This approach allows for evaluation of the relationship between these event probabilities and model parameters, which we refer to as arrhythmia sensitivity analysis.

We apply arrhythmia sensitivity analysis by analyzing the arrhythmogenic potential of mutations in the alpha subunit KCNQ1 of the slow-activating delayed rectifier current I_Ks_ that underlies Long QT Syndrome type 1 (LQTS1) [11]. Studies show that mutant I_Ks_ in LQTS1 contributes to generation of EADs, which may then lead to Torsades de Pointes at the level of the whole heart [12, 13]. Jons et al. described electrophysiological changes associated with 17 different LQTS1 mutations by experimentally measuring altered values of five I_Ks_-related parameters for each mutation [6]. We use the LRM to estimate and analyze how these measured changes in I_Ks_ parameters, and their experimentally observed uncertainties, impact P(EAD) for each of these LQTS1 mutations.

Action potential duration (APD) prolongation and QT-interval prolongation have been widely used as surrogate indicators for increased arrhythmogenic potential [14–17]. However, the relationship between these surrogate features and the likelihood of arrhythmia is indirect and sometimes at odds with experimental observations. For example, ranolazine leads to QT prolongation (a surrogate proarrhythmic indicator) in multiple clinical trials [18, 19] but has demonstrated anti-arrhythmic effects [20]. Recently, novel approaches for computational prediction of arrhythmogenic potential have emerged [21–24]. These studies utilize the abovementioned surrogates for arrhythmia to perform risk prediction. Rather than relying upon these indirect indicators of arrhythmic potential, in this work we develop model-based methods for directly estimating the probability of occurrence of the actual cellular events that trigger arrhythmias, EADs. Using the LQTS1 electrophysiological data of Jons et al. [6], we directly estimate P(EAD) for each LQTS1 mutation [25]. We compare the model estimated P(EAD) as an arrhythmia risk metric to those obtained using two other recently published methods. In the first method, Kernik et al. leveraged a population model of induced pluripotent stem cell derived cardiomyocytes (iPSC-CMs) to develop an approach for LQTS1 arrhythmia risk prediction[26]. In this approach, for each mutation, the parameters of the wild-type I_Ks_ in their control population model are modified using the distribution of parameters measured in mutant channels. For each mutation, the percentage of individual models in the population that satisfy all three of the following conditions is taken as the risk prediction metric: exhibit 1) > 4% increase of APD_90_, 2) ≥ 4% increase of beat-to-beat variation and 3) ≥ 4% increase of APD triangulation (APD_90_-APD_30_) [23]. The larger this percentage is, the greater the risk associated with the mutation. In the second method, Hoefen et al. used transmural repolarization prolongation (TRP) calculated using a 1-dimensional myocyte fiber model as an arrhythmic risk prediction metric [24]. Larger values of TRP are associated with greater risk.

We show that the relative risk (i.e., the rank ordered risk across the 17 mutations) estimated for the 17 LQTS1 mutations characterized by Jons et al. [6] are very similar across these three metrics. We also compare relative risk using these metrics to the relative clinical risk assessed using the 30-year cardiac event rate for each mutation [6]. All three risk metrics yield consistent estimates of the relative risk for the 17 mutations. In addition, the metrics are consistent in their identification of a subset of mutations whose predicted relative risk agree with, and another subset whose relative risks differ substantially from the measured relative clinical risk. A possible explanation for this discrepancy is that experimental characterization of these LQTS1 mutations is incomplete in functionally important ways.

## Results

### Modeling the Role of I_Ks_ on Probability of EADs

LQTS1 arises from mutations in KCNQ1, the alpha subunit of the I_Ks_ channel [11]. The symptoms of LQTS1 include syncope, cardiac arrest, and sudden death [27]. Jons et al. experimentally characterized the electrophysiological properties of I_Ks_ expressed in Xenopus oocytes for 17 LQTS1 mutations and reported the clinically measured 30-year cardiac event rate for each mutation [6]. The 30-year cardiac event rate is defined as the percentage of people within a population of carriers of the mutant who experienced syncope, cardiac arrest, or sudden death within the first 30 years of life. The outcomes of syncope, aborted cardiac arrest, and sudden cardiac death are defined as cardiac events.

An LRM was formulated to capture the quantitative relationship between P(EAD) and the 5 I_Ks_-related parameters measured experimentally (activation time constant scaling factor τ_+__sf; deactivation time constant scaling factor τ_-__sf; shift of the half maximal activation voltage ΔV_1/2_; scaling factor for voltage-dependent activation curve slope k_sf, and maximal conductance scaling factor G_Ks__sf). These 5 parameters were chosen because they have been experimentally measured for a number of different LQTS1 mutations [6]. Fig 1A shows examples of APs computed using the Walker et al. model [9] without (blue) and with (red) EADs. Because of the stochastic nature of the myocyte model, for the same parameter value, realizations obtained with different random number seeds may generate an AP with or without EADs. We defined a biophysically meaningful region of interest over which model parameters are varied and within which the LRM is built (S4 Table). Fig 1B demonstrates the relationship between parameters within this region of interest and P(EAD), where <u>B</u>^T^<u>P</u> (the argument of the logistic regression function) is the weighted (<u>B</u>) summation of LRM features (<u>P</u>). The nonlinear logistic relationship defines three domains of interest: transition domain, lower plateau domain, and upper plateau domain. The transition domain is a region in which P(EAD) is a sensitive function of <u>B</u>^T^<u>P</u>, corresponding to the steep part of the curve in Fig 1B. The lower plateau domain and upper plateau domain are regions where P(EAD) is relatively insensitive to parameter variation, with P(EAD) = ~0 in the lower plateau domain and P(EAD) = ~1 in the upper plateau domain. We sampled 200 parameter sets and ran 100 realizations for each set to train the LRM (see Methods). Once determined, the LRM enables direct calculation of P(EAD) for any parameter set without the need to simulate the underlying complex myocyte model. Fig 1B shows the fidelity with which the LRM model (blue line) reproduces P(EAD) estimated from simulations using the full myocyte model (red points) for the training data set (200 parameter sets) as a function of <u>B</u>^T^<u>P</u>. Detailed modeling methods and performance metrics are described in Methods and S1 Text. Fig 1C shows the LRM-predicted P(EAD) versus P(EAD) computed from the myocyte model simulations. The LRM performs well in predicting P(EAD) (R^2^ = 0.971 for the 96 parameter sets inside the transition domain, mean prediction error of 0.022 ± 0.036 (see S1 Text and Eq S1)), despite the highly complex nonlinear properties of the myocyte model. To test the ability of the LRM to generalize, Figs 1D-E compare the LRM predicted P(EAD) to that computed using the myocyte model for test data consisting of 100 independent parameter sets. The LRM performs well on the test set (R^2^ = 0.933 for the 27 parameter sets within the transition domain, mean prediction error = 0.019 ± 0.043).

**Fig 1.**
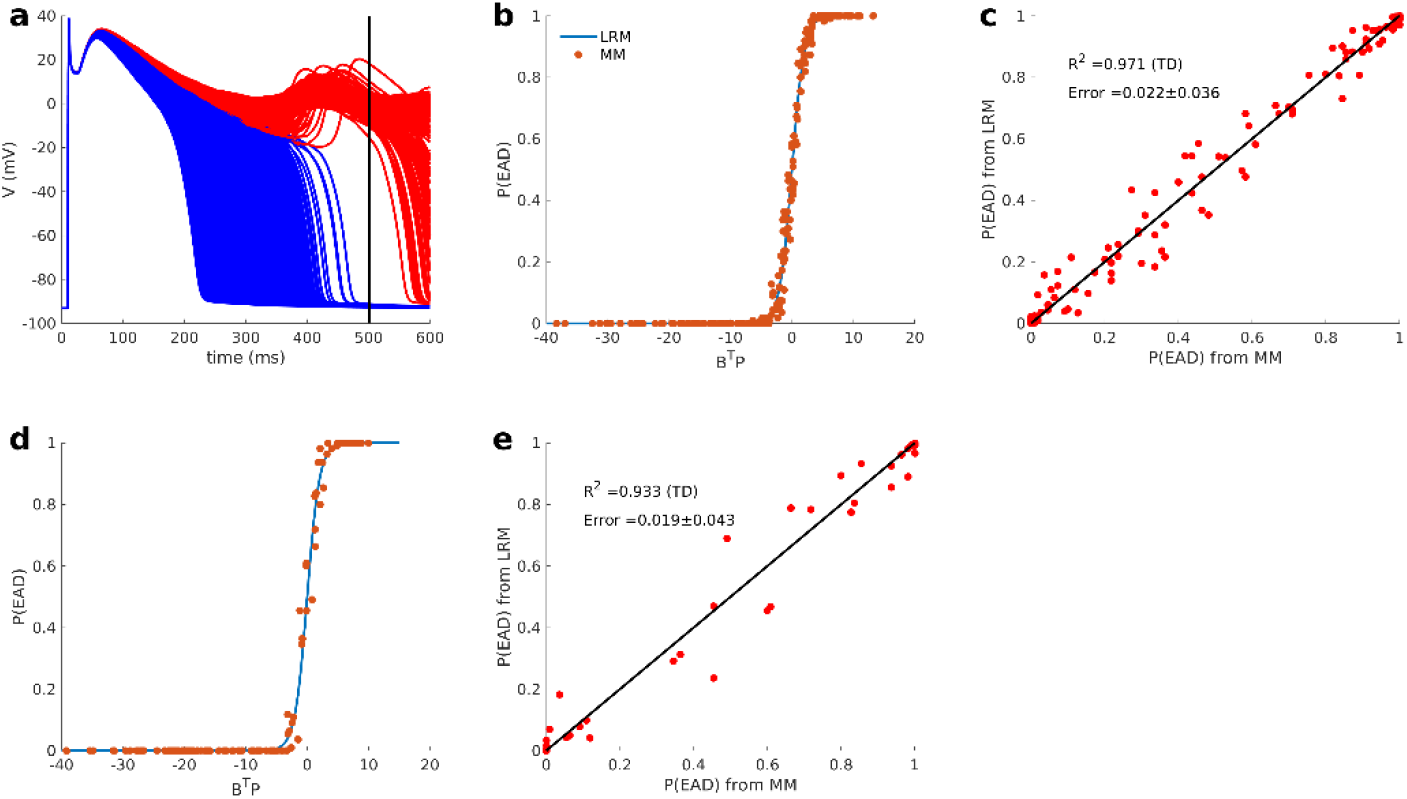
EAD study. (A) Example APs from myocyte model [9] simulations. Representative APs without EADs (blue) and with EADs (red) are shown. For each realization, an EAD is considered to have occurred if the membrane voltage at 500 ms (black vertical line) following the time of stimulus current is greater than −40 mV. (B) Comparison between LRM-predicted P(EAD) (blue) and P(EAD) computed from simulations using the myocyte model (red) as a function of <u>B</u>^T^<u>P</u> (weighted summation of LRM features) for the training set. (C) LRM-predicted P(EAD) (y-axis) vs corresponding P(EAD) estimated using the myocyte model (x-axis) for the training set. (D) and (E) show the same analyses described in (B) and (C), respectively, but for the independent test set. R^2^ is calculated only from parameter sets within the transition domain, LRM: logistic regression model, MM: myocyte model

The LRM for P(EAD) was formulated with a total of 16 features, 5 being the linear features (original parameters of I_Ks_) τ_+__sf, τ_-__sf, ΔV_1/2_, k_sf, and G_Ks__sf, and 11 being quadratic features (derived from the original I_Ks_ parameters) τ_+__sf*G_Ks__sf, ΔV_1/2_*G_Ks__sf, τ_+__sf^2^, τ_-__sf*G_Ks__sf, k_sf^2^, k_sf*ΔV_1/2_, τ_-__sf^2^, τ_+__sf*ΔV_1/2_, G_Ks__sf*k_sf, k_sf* τ_-__sf and τ_-__sf*ΔV_1/2_. Table 1 shows the LRM weights for each feature ranked by feature importance. In order to ensure interpretability of feature importance from the LRM weights, parameters were scaled to a range from 0 to 1. The data of Table 1 shows that I_Ks_ conductance (G_Ks__sf) is the most important, and this is followed in rank-order by τ_+__sf*G_Ks__sf, k_sf, and ΔV_1/2_*G_Ks__sf.

**Table 1.** EAD features ranked by relative importance.

| Ranking | Feature | Weight ± SE from LRM |
| --- | --- | --- |
| 1 | $G_{Ks\_sf}$ | $-63.89 \pm 1.86$ |
| 2 | $\tau_{+\_sf} * G_{Ks\_sf}$ | $34.35 \pm 2.07$ |
| 3 | $k\_sf$ | $14.57 \pm 0.65$ |
| 4 | $\Delta V_{1/2} * G_{Ks\_sf}$ | $12.83 \pm 1.61$ |
| 5 | $\tau_{+\_sf}$ | $11.90 \pm 0.59$ |
| 6 | $\tau_{+\_sf}^2$ | $-10.27 \pm 0.73$ |
| 7 | $\tau_{-\_sf} * G_{Ks\_sf}$ | $8.54 \pm 1.11$ |
| 8 | $k\_sf^2$ | $-7.67 \pm 0.41$ |
| 9 | $k\_sf * \Delta V_{1/2}$ | $-6.31 \pm 0.43$ |
| 10 | $\Delta V_{1/2}$ | $-6.05 \pm 0.51$ |
| 11 | $\tau_{-sf}^2$ | $-5.29 \pm 0.36$ |
| 12 | $\tau_{+sf} * \Delta V_{1/2}$ | $-4.66 \pm 0.69$ |
| 13 | $G_{Ks\_sf} * k_{sf}$ | $-4.00 \pm 1.11$ |
| 14 | $\tau_{-sf}$ | $3.09 \pm 0.62$ |
| 15 | $k_{sf} * \tau_{-sf}$ | $2.07 \pm 0.37$ |
| 16 | $\tau_{-sf} * \Delta V_{1/2}$ | $-1.26 \pm 0.37$ |

### Effect of LQTS1 Mutations on EAD Probability

For each mutation, 5 I_Ks_-related functional parameters (τ_+__sf, τ_-__sf, ΔV_1/2_, k_sf, and G_Ks__sf) were measured experimentally [6]. Measurements were obtained from no less than 16 oocytes for each mutation. Jons et al. quantified the experimental uncertainty in the measurement of each I_Ks_ parameter for each mutation by providing a mean (μ) and standard error for each of the five experimentally measured parameters. We derived the standard deviation (σ) from the reported standard error, which is summarized in S1 Table.

The LRM is used to investigate the effects of experimental uncertainty in parameter estimates on P(EAD) for each mutation. Parameter sets are constructed by choosing each of the five parameters as independent normally distributed random variables with mean μ and standard deviation σ measured experimentally. For each of these randomly generated parameter sets, the LRM maps these parameters to P(EAD). Therefore, the value of P(EAD) predicted using the LRM is itself a random variable with its own distribution. We estimate this distribution using Monte Carlo methods by choosing 10^6^ normally distributed parameter sets for each mutation (see details in Methods). Parameter set distributions estimated in this way reveal the impact of experimental uncertainties on estimates of P(EAD) dervied from arrhythmia sensitivity analysis.

Fig 2A-C show the distribution of P(EAD) for the three representative mutations *G314S*, *G168R*, and *Y315C*, with high, medium, and low average P(EAD), respectively. The P(EAD) distribution for *G314S* is narrow with a single peak at P(EAD) near one. This indicates that despite the experimental uncertainty (EU) in measurement of this parameter set, we can be quite certain that this mutation leads to EADs with high probability. However, the *G168R* and *Y315C* distributions are bimodal with peaks near zero and one. In these cases, parameter sets belong to two subgroups, one that almost always leads to EADs and another that seldom does. These bimodal distributions of P(EAD) can be approximated by a Bernoulli distribution with parameter p, where p = P(P(EAD) >0.5) for any particular parameter set and 1-p = P(P(EAD) <0.5). The parameter p, which can be approximated as the mean P(EAD), denoted P_m_(EAD), is shown by the red vertical line in Figs. 2B and C, where *G168R* has larger mean P(EAD) than *Y315C*.

**Fig 2.**
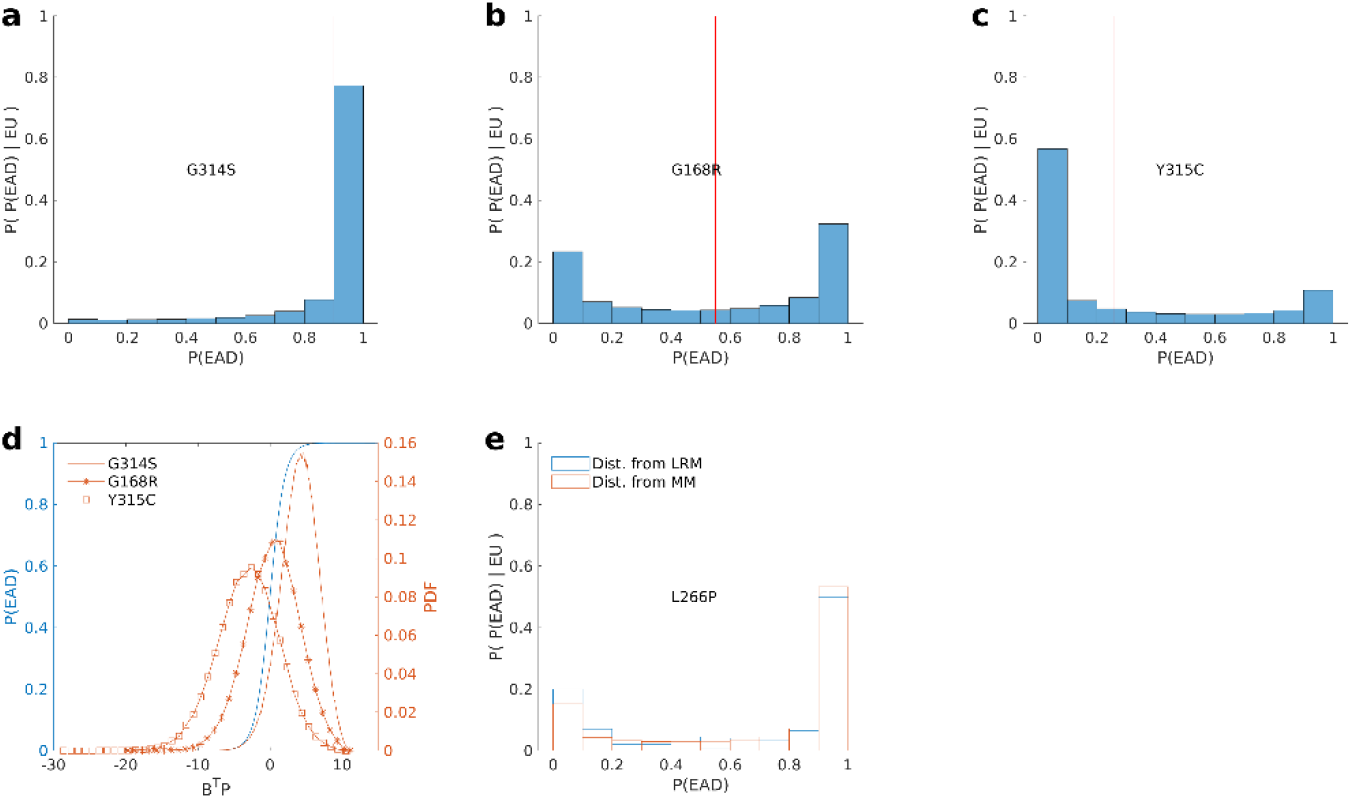
LQTS1 study. LRM-based estimates of P(EAD) distribution for mutations: (A) *G314S*, (B) *G168R*, and (C) *Y315C*. Red vertical lines in A-C represent the mean P(EAD) given the observed experimental uncertainty (EU) in each mutation-specific parameter set. This is denoted as P(P(EAD) | EU) on the ordinate. (D) PDFs of <u>B</u>^T^<u>P</u> (red) are shown for each mutation (*G314S*: solid line, *G168R*: star markers, and *Y315C*: square markers), where <u>B</u>^T^<u>P</u> is the weighted summation of 16 I_Ks_ features (5 parameters + 11 quadratic features) for each mutation. The logistic function curve (blue) relates <u>B</u>^T^<u>P</u> to P(EAD). (E) Comparison of P(EAD) distributions for *L266P* computed using the LRM (blue) and myocyte model (red).

To explain the observed P(EAD) distributions, the probability density function (PDF) of <u>B</u>^T^<u>P</u> (red) and the model logistic function (blue) that relates <u>B</u>^T^<u>P</u> to P(EAD) is shown in Fig 2D. Recall that <u>B</u>^T^<u>P</u> is a function of 5 I_Ks_-related parameters which appears in the LRM. Therefore, probability density functions (PDFs; red) of <u>B</u>^T^<u>P</u> can be derived using the Monte Carlo method with parameter distributions matching the experimental uncertanties measured by Jons et al. [6]. For mutation *G314S*, the PDF (squares) is narrow with mean positioned within the upper plateau domain and part of the transition domain of the model logistic function. This results in a P(EAD) distribution that is unimodal and close to 1. For the *G168R* and *Y315C*, PDFs are broader and span all domains (i.e. lower plateau domain, transition domain, and upper plateau domain) of the model logistic function, thus resulting in a P(EAD) distribution that is bimodal.

To validate this use of the LRM in uncertainty analyses, we performed Monte Carlo simulations of the myocyte model with 200 randomly generated sample parameter sets for the *L266P* mutation (200 realizations were run for each set). Fig 2E shows that the distribution of P(EAD) predicted by the LRM faithfully reproduces that generated from the myocyte model. The total computation time required for all myocyte model simulations was ~17,000 CPU hours whereas the LRM model required less than 1 CPU second.

To further delineate the effects of EU in the measured properties of I_Ks_ for each mutation, we evaluated the contribution of each individual parameter to the uncertainty of P(EAD) for each mutation. To do this, only a single parameter is varied while others are assigned a constant value equal to their experimentally measured mean value, and this process is repeated for each parameter set. In S2A Fig, PDFs (red curves) are shown for *G314S* and *Y315C* arising from variations in τ_+__sf based on its EU with all other parameter sets held constant. S2B-E Figs show the same analyses for τ_-__sf, ΔV_1/2_, k_sf, and G_Ks__sf. Note that the PDFs in Fig 2D, where all parameters vary simultaneously, are nearly identical to those of S2E Fig, indicating that G_Ks_ uncertainty plays a dominant role in determining overall uncertainty of the P(EAD) across all 5 I_Ks_ parameters for all mutations.

### LQTS1 mutation arrthymic risk prediction

Kernik et al. designed an arrhythmia risk prediction metric that is calculated using a population model of myocytes (iPSC-CMs) [28]. In this approach, a population model is built for each mutation by replacing the wild-type I_Ks_-related parameters in their control population model with those chosen from a distribution of mutation parameters. The risk prediction metric is taken as the percentage of model myocytes in the population model for each mutation satisfying all 3 of the following conditions: 1) > 4% increase of APD_90_, 2) ≥ 4% increase of beat-to-beat APD_90_ variation, and 3) ≥ 4% increase of APD triangulation (APD_90_-APD_30_)[23]. We refer to this metric as the AP morphology metric and calculated it for the 17 LQTS1 mutations using this population myocyte model.

Hoefen et al. used a modified Flaim-Giles-McCulloch [29] 1D fiber model to estimate transmural repolarization prolongation (TRP) for these 17 mutations [15]. We denote this metric as the TRP metric. Values of this metric for the 17 LQTS1 mutations were taken from Fig 1C of Hoefen et al. [15]. Further details are given in Methods.

S2 Table shows prediction of three model-based risk metrics for 17 mutations. S3 Fig shows that P_m_(EAD) is well aligned with the AP morphology metric (R^2^ = 0.93, S3A Fig) and the TRP metric (R^2^=0.80, S3B Fig). To compare model-based risk metrics with clinical risk, we determined the relative risk ranking (r) for each mutation (with rank of 1 denoting the lowest risk and a rank of 17 denoting the highest risk amongst the 17 mutations). The relative risk ranking predicted using a model-based metric is denoted as r_pred_ and the relative risk ranking based on clinical risk is denoted as r_clin_. For example, *A341V* has r_clin_=17 (highest clinical risk) and r_pred_ = 6, 8, or 5 (intermediate predicted risk) for P_m_(EAD), AP morphology metric, and TRP metric, respectively. S2 Table shows r_pred_ of the three different risk metrics and r_clin_ for the 17 mutations. We then calculated the difference (Δr = r_pred_ - r_clin_) between the relative risk ranking predicted using each of the three model-based metrics and the clinical ranking. If Δr is negative, then r_pred_ underestimates arrhythmic risk, and vice versa. The greater the difference in ranking, the greater is the under-(Δr < 0) or over-(Δr > 0) prediction. Fig 3 shows Δr for all 3 metrics for the 17 mutations. In Fig 3, mutations are ordered from those with the greatest underprediction of risk to those with the greatest overprediction of risk. The three metrics yield similar results for all mutations, consistent with the relationships among the metrics shown in S3 Fig. Results show that *A341V* and *R494Q* are the two mutations with the most severely underpredicted risk, while *G168R* and *W305S* are the two mutations with the most overpredicted risk. The r_pred_ for mutations S225L, R243C, V254M, D611Y and G314S are in close agreement (−3 ≤ Δr ≤ 3) with their respective r_clin_.

**Fig 3.**
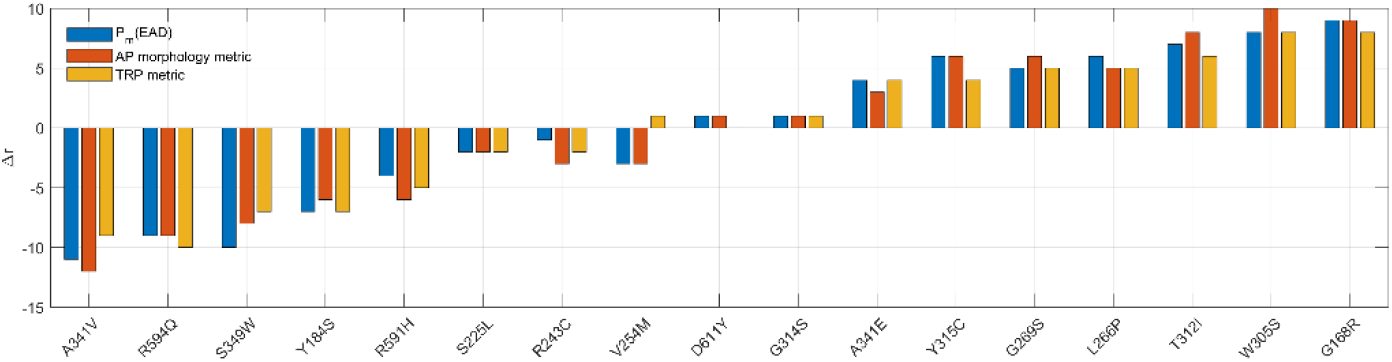
Relative risk ranking comparison across 3 model-based risk metrics. The difference of relative risk ranking (Δr = r_pred_ - r_clin_) between that predicted using the model-based metric (r_pred_) and that based on clinical risk (r_clin_) for all 17 mutations. For each mutation, there are three r_pred_ values corresponding to each of the three model-based risk metrics. An average r_pred_, and hence an average Δr, is calculated for each mutation and determines their ordering (from most negative to most positive Δr)

In the study of Jons et al. [15], different numbers of patients exhibited each mutation. The number of patients evaluated with a given mutation may therefore limit the statistical significance of the risk associated with that mutation. Therefore, we performed a pairwise Fisher exact test for each pair of mutations (S4 Fig) to determine those pairwise differences in risk that are statistically significant in the Jons et al. study. 42 out of 136 mutation pairs with p<0.05 were found to be significant pairs. Three approaches (P_m_(EAD) in Fig 4A, AP morphology metric in Fig 4B, and TRP metric in Fig 4C) are used to rank order the risk for significant pairs and to compare them with the rank order of clinical risk. An entry of 1 indicates that the predicted rank order of the mutation pair is consistent with the clinical rank order for this pair, whereas an entry of 0 means the predicted rank order is reversed and opposite to that observed clinically. For example, for *V254M-W305S* pair, *V254M* has greater clinical risk than *W305S*, thus the clinical rank order is *V254M* > *W305S*. Given the predicted risk from P_m_(EAD) of *V254M*(P_m_(EAD) = 0.742) and *W305S* (P_m_(EAD) = 0.184), the predicted rank order is *V254M* > *W305S*, consistent with the clinical rank order. Thus, in Fig 4A, the element in the lattice for the *V254M-W305S* pair is set to 1. In contrast, based on the prediction by AP morphology metric, the predicted rank order is *V254M* < *W305S*. Thus, in Fig 4B, the element in the lattice for the *V254M*-*W305S* pair is set to 0.

**Fig 4.**
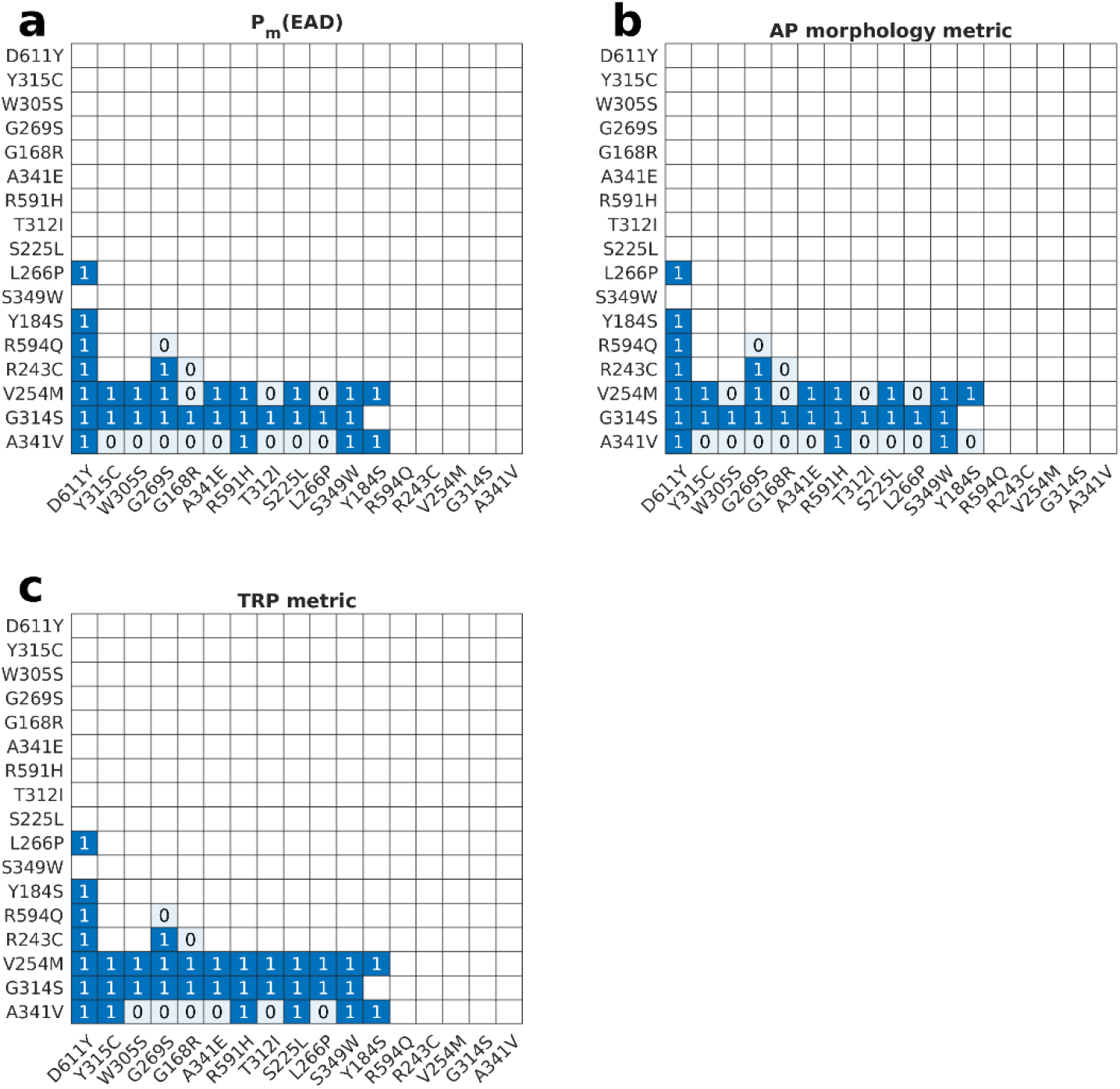
Performance of pairwise prediction in significant mutation pairs. Three approaches, (A) P_m_(EAD), (B) AP morphology, and (C) TRP, are used to predict the risk rank order of significant mutation pairs. Mutations are ordered by clinical risk ranking from lowest to highest (left to right on the x-axis and top to bottom on the y-axis). A table value of 1 indicates that, for this pair, the ranking of r_pred_ values is consistent with that of r_clin_ values and a table value of 0 means that predicted rank is inconsistent with the clinical rank. Since significant pairs are shown only in the lower triangle of the heatmap, for any pair, the y-axis mutation always has a greater clinical risk than the x-axis mutation.

To quantify performance of each metric, the accuracy of pairwise relative risk prediction is measured by calculating the percentage of mutation pairs for which the predicted relative risk matched the clinical relative risk. The set of mutation pairs was divided into two subgroups, one in which all pairs have a statistically significant clinical risk difference (p <0.05 in the Fisher exact test), and its complement. The accuracy for each metric is summarized in Table two across all mutation pairs, as well as separately within these two subgroups. For all metrics, the accuracy within the subgroup of significant pairs is higher than the accuracies across all pairs and or within the subgroup of non-significant pairs. The TRP metric has the highest accuracy in both significant pairs and all pairs. Despite this, there remain 37% of mutation pairs whose relative risk the TRP metric is unable to predict correctly. The detailed pairwise prediction on all pairs is shown in S5 Fig. In Fig 4, all metrics performed poorly on *A341V*-associated pairs, which is consistent with the conclusion from Fig 3 that relative risk associated with mutation *A341V* is underestimated.

## Discussion

### Logistic regression model on probability of EAD

Using our previously developed method [10], we built a logistic regression model that maps 5 I_Ks_-related parameters to P(EAD). Fig 1E shows that P(EAD) predicted using the LRM agrees well with that estimated from simulations obtained using the Walker model. The computational efficiency of the LRM approach is what makes arrhythmia sensitivity analysis for experimentally determined parameter set distributions possible. In Fig 2E, calculating the *L226P* P(EAD) distribution using the Walker myocyte model (200 parameter sets and 200 realizations for each set) required ~16,000 CPU hours whereas the LRM required ~0.5 CPU second to simulate the 10^6^ parameter sets.

### Arrhyhmia sensitivity analysis

LRMs enable us to quantitatively explore the relationship between parameters and P(EAD). G_Ks_ is the most important parameter of I_Ks_ with regard to P(EAD) because in the ranking of feature importance, it is associated with the first, second, and fourth features in Table 1. To analyze the relationship between parameters and P(EAD), we vary one parameter while fixing the other four parameters at their wild type values. S1A Fig shows that decreasing G_Ks_ results in P(EAD) going from 0 to 1. This result follows from the fact that I_Ks_ is a repolarizing current. With the decrease of conductance of I_Ks_, the action potential becomes more prone to depolarizing events, thus leading to higher likelihood of EAD occurrence [30]. However, with G_Ks_ at its wild-type value, varying any other parameter has no effect on P(EAD), which is always 0. Furthermore, when G_Ks_ is reduced to half of its wild-type value and other parameters are varied (τ_+__sf (S1B Fig), τ_-__sf (S1C Fig), ΔV_1/2_ (S1D Fig), and k_sf (S1E Fig)), P(EAD) is positively related to τ_+__sf, ΔV_1/2_, and k_sf. This indicates that a reduction of G_Ks_ not only increases P(EAD), but also increases the sensitivity of the P(EAD) to variation of other parameters.

### Uncertainty analysis on LQTS1 mutations

In Fig 2B and 2C, a bimodal distribution of P(EAD) for mutations *G168R* and *Y315C* is observed. Fig 2D shows that this arises as an interplay between the PDF of <u>B</u>^T^<u>P</u> (when parameters <u>P</u> are modeled as normally distributed random variables, <u>B</u>^T^<u>P</u> is also a random variable with a normal distribution) and the model logistic function. Similar bimodal distributions were observed in our previous study of the probability of ectopic beats caused by delayed afterdepolarizations [10]. Given the bimodal shape of the P(EAD) distribution, small perturbations of the 5 I_Ks_-related parameters result in model cells that either exhibit normal APs exclusively, or APs with EADs exclusively.

In S2 Fig, we evaluated the contribution of each individual parameter to the uncertainty of P(EAD) for *G168R* and *Y315C*. The PDF of <u>B</u>^T^<u>P</u> in S2E Fig arising from variability in G_Ks__sf alone nearly matches the PDF in Fig 2D, where variabilities of all parameters are considered, indicating that G_Ks__sf uncertainty alone provides enough information to approximately reconstruct the P(EAD) distribution. This result indicates that in LQTS1, I_Ks_ expression variability is more impactful than variations in its kinetic or voltage-dependent properties with respect to the uncertainty of P(EAD). A similar conclusion has been made in LQTS type 2 (LQTS2). Anderson et al. showed that protein trafficking (which ultimately impacts cell-surface expression), rather than gating or permeation (intrinsic functional properties of the channel), is the dominant mechanism for loss of Kv11.1 channel function in LQTS2 [31].

### The comparison of arrhythmia risk prediction approaches

Action potential duration (APD) and QT prolongation have been widely used as surrogates, both in basic research and in clinical settings, as indicators of disease [14, 15] and drug-related arrhythmogenic effects [16, 17]. However, there are examples which have shown the dissociation of these two surrogates with risk of arrhythmia. For example, alfuzosin prolongs APD and QT interval but is not linked to occurrence of Torsades de Pointes [32]. Ranolazine leads to QT prolongation (surrogate proarrhythmic indicator) in multiple clinical trials [18, 19] but has demonstrated anti-arrhythmic effects [20]. Computational methods of arrhythmia risk prediction have recently emerged[15, 21–23]. These studies focus on simulating the abovementioned surrogates and other indirect surrogates to predict arrhythmia risk. Unlike these indirect surrogates, the EAD is a fundamental cellular arrhythmia event. The stochastic myocyte model, in principle, can be used to estimate P(EAD) but doing so is unrealistic because of its high computational burden. Such estimates become possible with our highly simplified LRM approach.

We not only examined P_m_(EAD) as a risk metric, we also contrasted its performance with two other published metrics: 1) AP morphology metric (a metric based on AP morphology over a population of myocytes) [33]; and 2) TRP metric (transmural repolarization prolongation within a 1D cardio fiber model) [24]. S3 Fig shows that all three approaches yield very similar predictions of relative risk. Fig 3 shows that the predicted relative risk is consistent across all three model-based risk metrics. Note that calculation of each metric is done using a different myocyte model. The Walker stochastic model is used for P_m_(EAD). The Kernik iPSC-CM population model [28] is used to calculate the AP morphology metric. A modified Flaim-Giles-McCulloch [29] 1D fiber model is used for the TRP metric. Thus, results are consistent across risk metrics and across the different computational models of the cardiac myocyte on which they are based. Table 2 shows that the TRP metric has the highest accuracy (0.63) in pairwise mutation relative risk prediction.

**Table 2.**
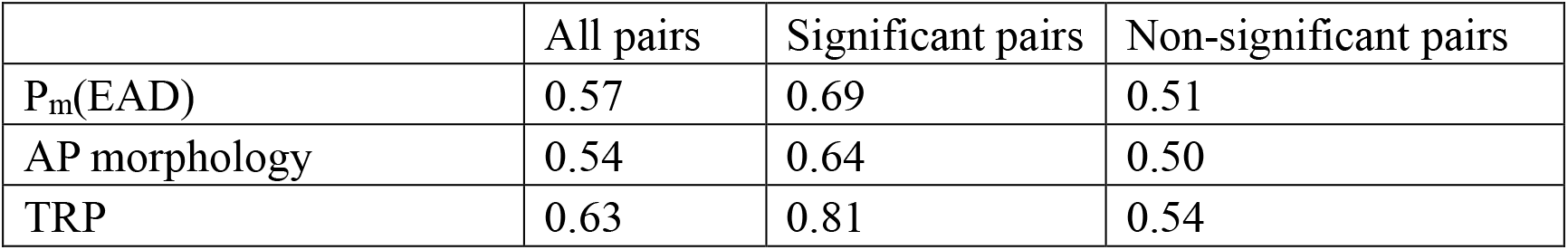
Mutation pairwise prediction accuracy

|  | All pairs | Significant pairs | Non-significant pairs |
| --- | --- | --- | --- |
| $P_m(\text{EAD})$ | 0.57 | 0.69 | 0.51 |
| AP morphology | 0.54 | 0.64 | 0.50 |
| TRP | 0.63 | 0.81 | 0.54 |

To test the generalizability of the P_m_(EAD) metric, we also assessed P_m_(EAD) using another stochastic myocyte model, specifically the model of Greenstein et al [25]. A major difference between these two models is that the Walker model is a spatial model of the myocyte, whereas the earlier Greenstein model is a stochastic, non-spatial, compartmental model. We denote the P_m_(EAD) calculated from the Greenstein model as P_m_(EAD)_G_. S3C Fig shows that P_m_(EAD)_G_ is well aligned with the P_m_(EAD) (R^2^ = 0.91) across these two different stochastic models.

qNet is another well-studied drug arrhythmic risk prediction metric based on a modified O’hara Rudy human myocyte model [34]. Detailed results using qNet are shown in S2 Table [34]. S3D, Fig shows that qNet is substantially different from P_m_(EAD) for the 17 mutations in this study (R^2^ = 0.07). In S6 Fig, the qNet metric yields a substantially different prediction of relative risk order compared with the three approaches described here. Parikh et al. performed a global sensitivity analysis on qNet and showed that qNet is insensitive to the perturbation of I_Ks_ current parameters[35]. For this reason, we chose not to focus on use of qNet for predicting the risk of LQTS1 mutations in this study.

Clearly, each of the three metrics studied make significant errors in predicting the relative risk of mutations. Nonetheless, these intrinsically different metrics based on estimates of P_m_(EAD), AP morphology across populations of myocyte models (the AP morphology metric), and a 1-D tissue-level model (the TRP metric) yield highly consistent predictions of relative risk across the mutations studied. This consistency across metrics and models suggests to us that for those mutations where risk is significantly under- or over-estimated, the fundamental properties of these mutations may be inadequately characterized by the five I_Ks_ parameters studied by Jons et al. [15].

We give a possible example. Heijman et al. found that in the LQTS1 mutation *A341V*, not only is I_Ks_ reduced, but that the increase of I_Ks_ observed during βAR-stimulation in wild-type myocytes is reduced. Given that ~ 88% of triggered cardiac events in LQTS1 patients occurs during exercise or emotional stress [36], the characteristics of I_Ks_ during βAR-stimulation are relevant to evaluating arrhythmia risk. In the P_m_(EAD) approach, our modeling incorporated βAR-stimulation effects. However, the degree of I_Ks_ upregulation was considered to be the same as that measured in wild type I_Ks_. When we incorporate the Heijman et al. results by reducing βAR-induced upregulation of I_Ks_, P_m_(EAD) in the *A341V* mutation increases from 0.17 to 0.50, and overall relative risk of this mutation within the total 17 mutations increases from rank 6 to rank 12. Mutation *A341V* has the highest clinical risk. Although there remain components of risk that are unaccounted for, incorporating βAR effects of this mutation does increase risk estimated by P_m_(EAD). Additionally, Heijman et al. also observed that another LQTS1 mutation *G589D* causes the loss of βAR-mediated I_Ks_ upregulation. This indicates that the suppression effect of I_Ks_ upregulation during βAR-stimulation may also occur on other LQTS1 mutations. These data suggest that the effects of βAR stimulation on I_Ks_ in LQTS1 mutations are an essential, yet lacking component for risk prediction.

Values of P(EAD), as calculated in this study, do not represent the actual probabilities of EAD-induced cardiac events given the fact that the electrotonic loads across coupled cells in tissue are not taken into consideration. The electrotonic load at the location of any cell in the heart counteracts effects of depolarizing current by shunting charge out of the cell across gap junctions, thereby reducing the likelihood of triggering events such as EADs. This load is reduced if neighboring cells depolarize more synchronously, and under this scenario a collection of cells may locally generate EADs that can successfully propagate in the heart [37]. Therefore, the probability of a propagating tissue-level cardiac event will be much smaller than P(EAD) predicted in an isolated cell model. However, we believe that P(EAD) is intrinsically related to the fundamental probability of tissue-level cardiac events because, given the same electrotonic load, the probability of cardiac events is clearly positively correlated with P(EAD) in a single myocyte, as these events arise from a population of EAD generating cells. Therefore, understanding how mutations and other cellular alterations impact the generation of cellular arrhythmias such as EADs is a fundamental step in understanding how they lead to clinical cardiac events.

In summary, we developed a LRM mapping 5 I_Ks_-related parameters to P(EAD). We used this model to investigate how variation of I_Ks_-related parameters affects P(EAD).

A reduction of G_Ks_ increases P(EAD) and increases its sensitivity to variation of other parameters. We also examined how experimental uncertainties of I_Ks_-related parameters affect model estimated distributions of P(EAD) and their interpretation. The experimental uncertainty of G_Ks_ is more impactful than uncertainties in its kinetic or voltage-dependent properties with respect to the uncertainty of P(EAD). Two other model-based risk metrics were studied, and results were compared to those obtained by calculating mean P(EAD). All approaches yield very similar predictions regarding the relative risk of 17 I_Ks_ mutations, and this result holds despite the fact that these risk metrics and the models to which they are applied are very different. These three model-based risk metrics yield similar prediction performance; however, they all fail to predict clinical risk for a significant number of the 17 studied LQTS1 mutations. This suggests that experimental characterization of a number of LQTS1 mutations may be incomplete.

## Methods

### Modified Walker Ventricular Myocyte Model

This study uses a modified version of the Walker et al. stochastic ventricular myocyte model [9] described in Jin et al. [10]. A subset of CRUs (5400 out of 25000) was used for all simulations to accelerate the computation. β-adrenergic stimulation is simulated to facilitate generation of cellular arrhythmias. This was done using the β-adrenergic stimulation protocol described in Walker et al. [9]. The model is available in Github (https://github.com/JHU-Winslow-Lab/EAD-paper-code.git).

### Long QT syndrome type 1 experimental data and IKs model

LQTS1 clinical and experimental electrophysiological data including 30-year cardiac event rate for 17 mutations are obtained from Jons et al. 2011 [6]. The 30-year cardiac event rate represents the percentage of mutation carriers that had their first cardiac event in their first 30 years of life. Cardiac events include syncope, aborted cardiac arrest, and sudden cardiac death. Clinical and experimental data from their study is summarized in S1 Table. Five I_Ks_-related parameters for each mutation are taken from from Jons et al. Tables 2 and S1. In order to incorporate these five I_Ks_-related parameters into our myocyte model, we also incorporated their I_Ks_ model into the Walker model. The I_Ks_ model is described by the following equations:

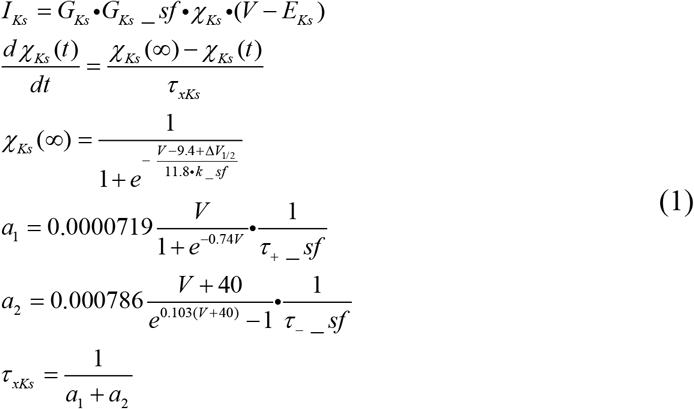

where τ_+__sf, τ_-__sf, ΔV_1/2_, k_sf, and G_Ks__sf are mutation-specific terms.

### Simulation protocol for early afterdepolarization modeling

In the EAD studies, initial states of model state variables were the same in each simulation. These values were obtained in diastole after 10 beats at 1 Hz pacing during the β-adrenergic stimulation protocol. Each model simulation had a duration of 600ms. At 10ms, a 1-ms 10000 pA inward current was injected to trigger an action potential.

### Logistic regression model

We recently developed a workflow to build a logistic regression model (LRM) on complex myocyte models [10]. It maps model parameters to the probability of a cellular arrhythmia event using the following Logistic equation:

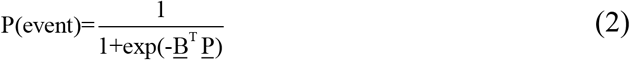

where <u>P</u> is a vector of features and <u>B</u> is a vector of weights. Features include model parameters and their products. See S1 Text for details description on the LRM method.

### Implementation of arrhythmia risk prediction metrics

Implementation details of P_m_(EAD)_G_, AP morphology metric, and qNet are described in S2 Text. Models and results are available in Github (https://github.com/JHU-Winslow-Lab/EAD-paper-code.git).

## Supporting information

**S1 Fig.**
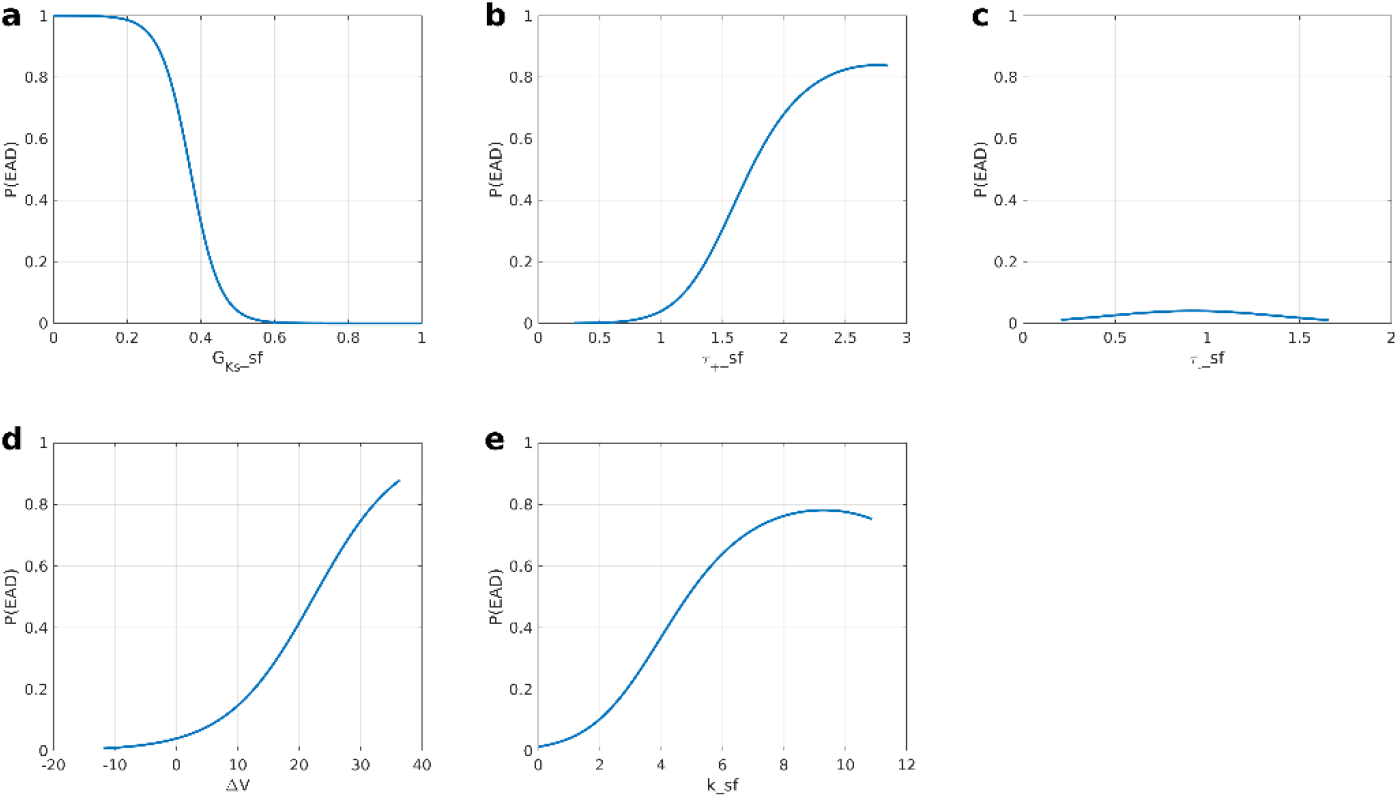
Relationship between 5 IKs-related parameters and P(EAD). (A) vary the G_Ks__sf while fixed other four parameters at wild type value (τ_+__sf = 1, τ_-__sf = 1, ΔV_1/2_ = 0, and k_sf = 1). Similarly, vary one parameters: τ_+__sf (B), τ_-__sf (C), ΔV_1/2_ (D) or k_sf (E) and fix other parameters at wild type except that G_Ks__sf = 0.5.

**S2 Fig.**
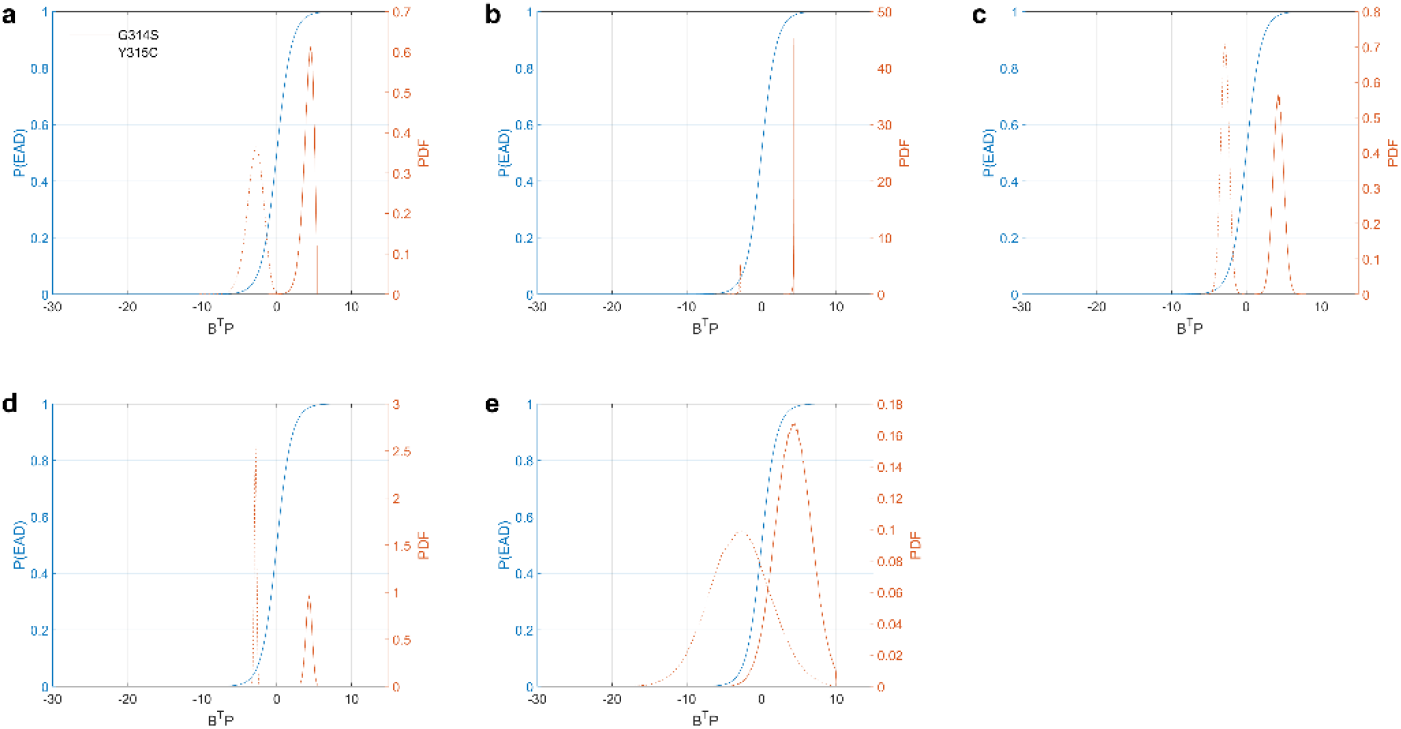
Uncertainty analysis for individual features underlying LQTS1 mutations. PDFs of <u>B</u>^T^<u>P</u> (red) are shown for two mutations (*G314S*: solid line and *Y315C*: square markers). <u>B</u>^T^<u>P</u> is the weighted summation of 16 I_Ks_ features (5 parameters +11 parameter-derived quadratic features) for each mutation. The MC curve (blue) relates <u>B</u>^T^<u>P</u> to P(EAD). Single feature uncertainty: (A) τ_+__sf, (B) τ_-__sf, (C) ΔV_1/2_, (D) k_sf, or (E) G_Ks__sf is sampled from an experimentally based distribution while other parameters are assumed fixed at their mean.

**S3 Fig.**
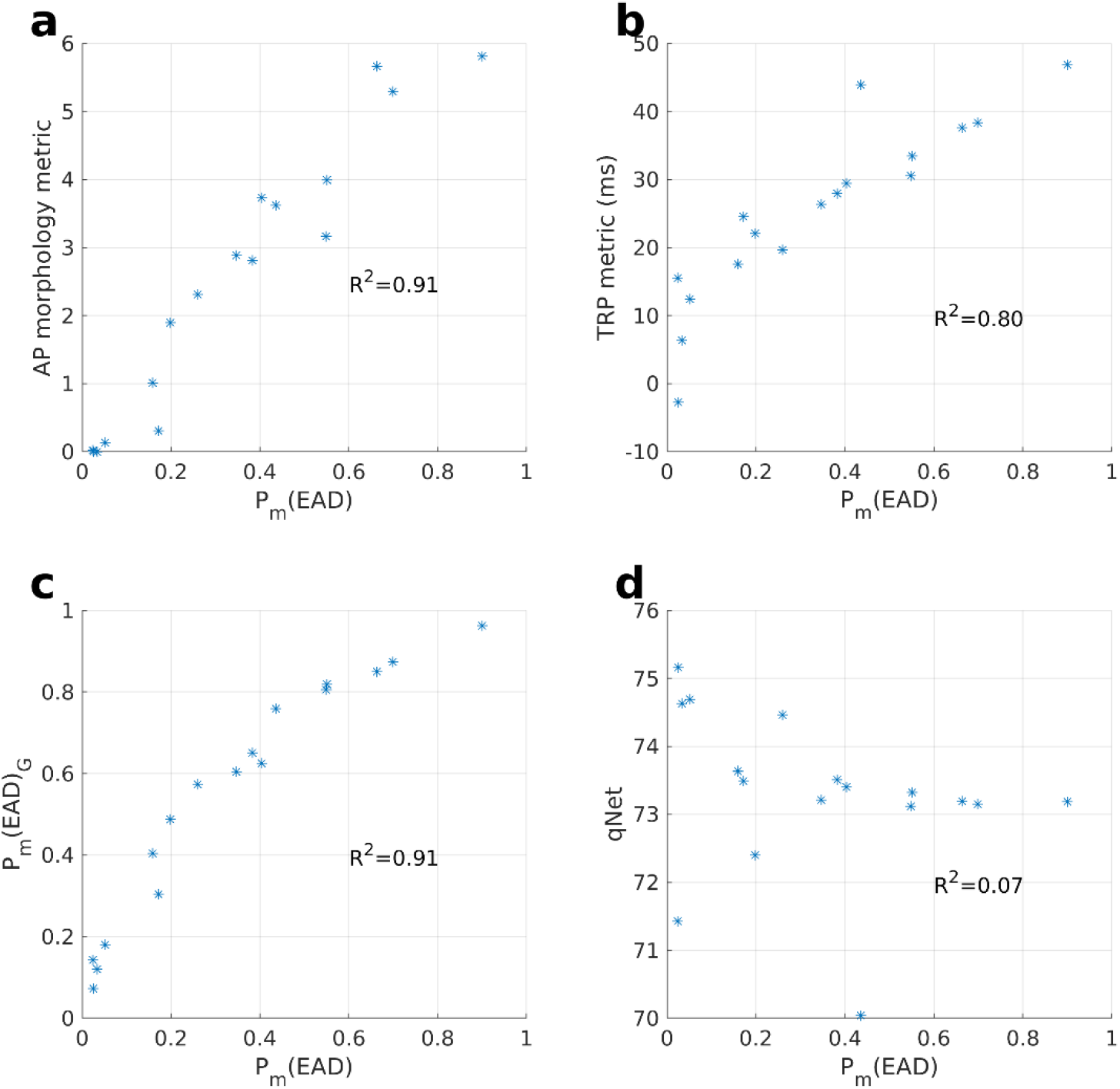
Comparison between model-based risk metrics. The comparison between P_m_(EAD) (at x axis) and other three metrics (at y axis) on 17 mutations: (A) AP morphology metric, (B) TRP metric, (C) P_m_(EAD)_G_ and (D) qNet. Linear regression between two compared approaches is performed and R^2^ is reported.

**S4 Fig.**
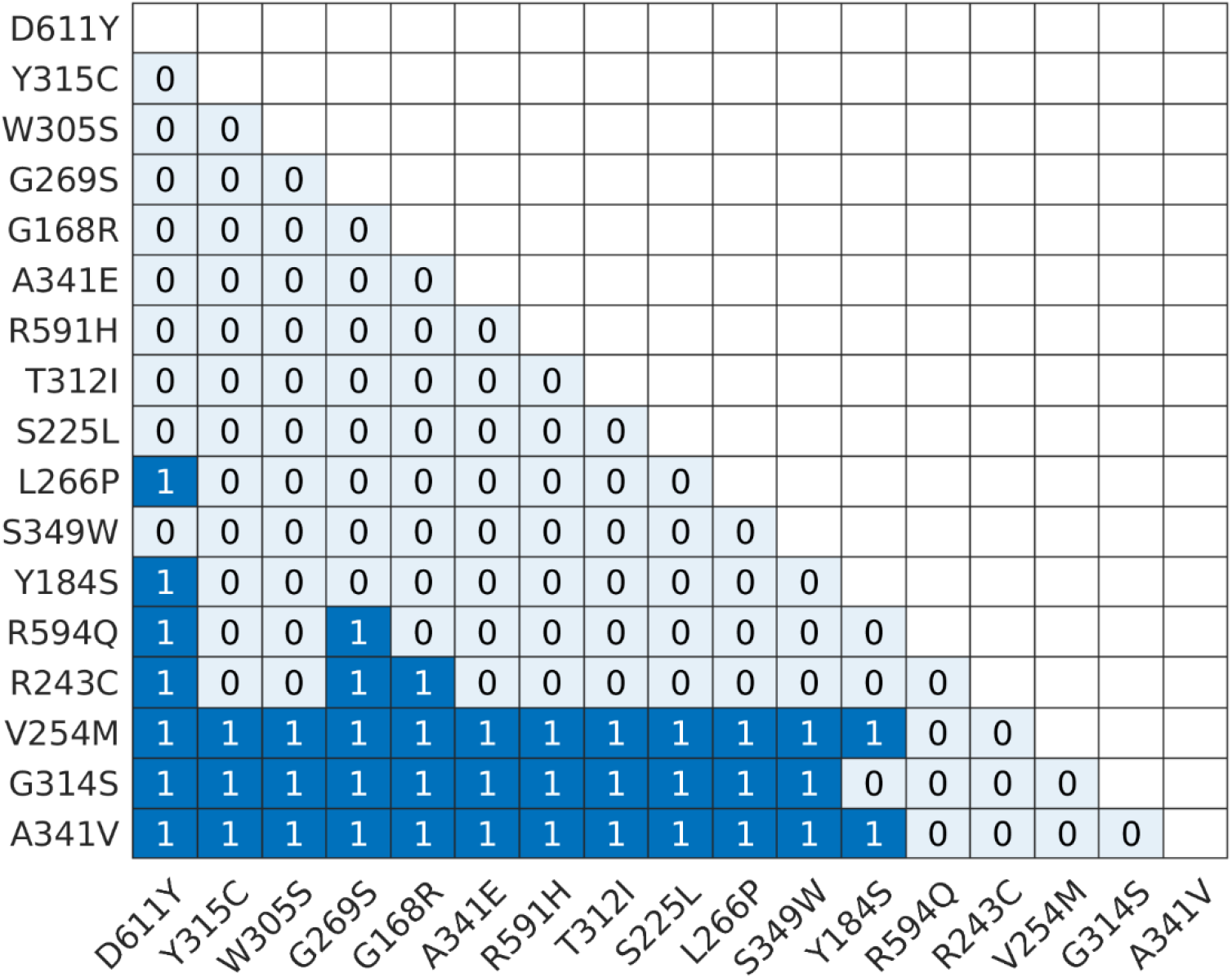
Fisher exact test on 17 mutations. The fisher exact test is performed on every mutation pairs to test if the difference of clinical risk (30-year cardiac event rate) is statistically different. 1 represents that the mutation pair has significant clinical risk difference. 0 means that clinical risk difference is not significant. On x axis, from left to right (and on y axis from top to bottom), mutations are ordered the clinical risk from least to most.

**S5 Fig.**
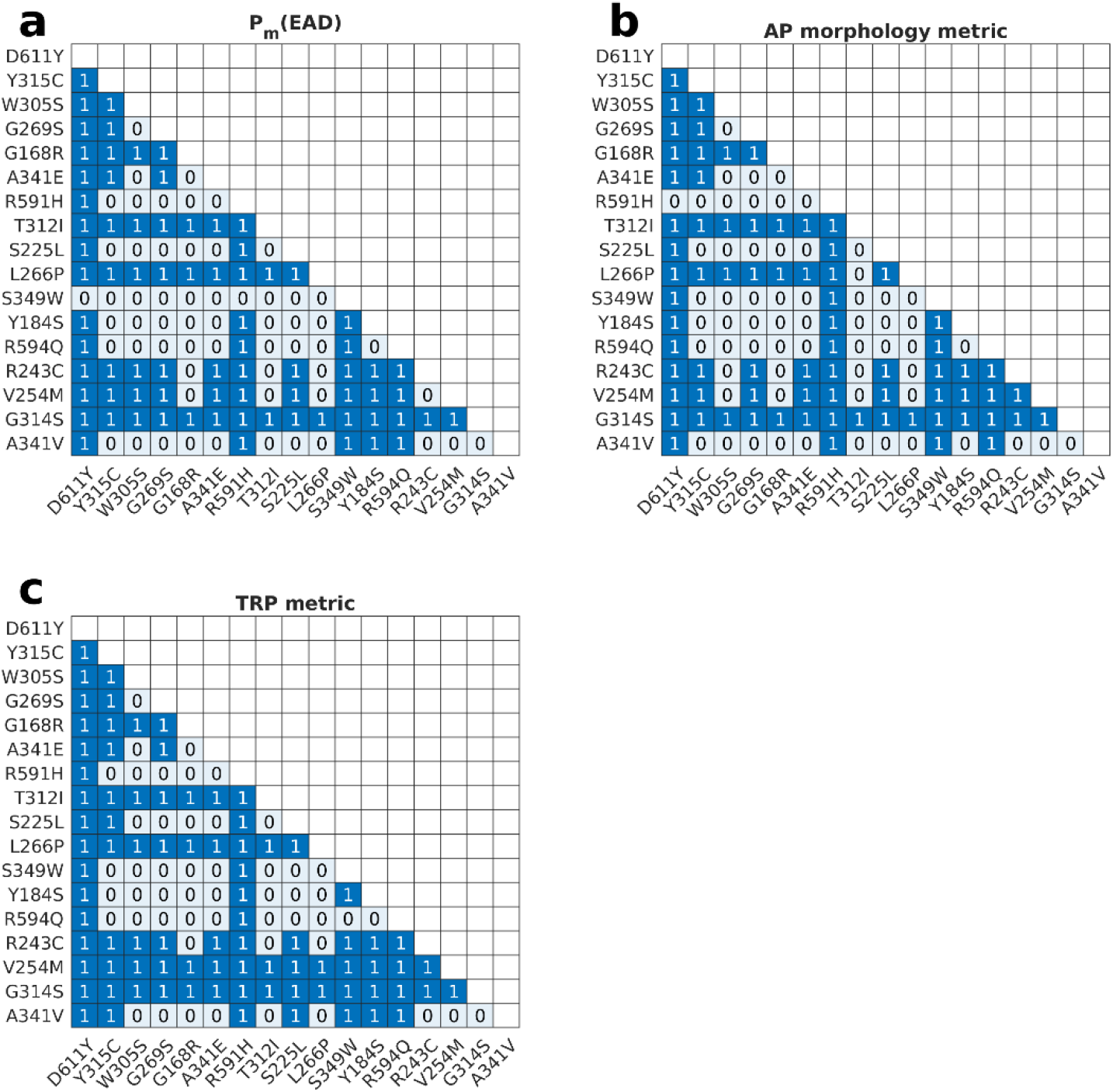
Pairwise prediction in all mutation pairs. Similar figure as Fig 4 except that all pairs are considered in this analysis.

**S6 Fig.**
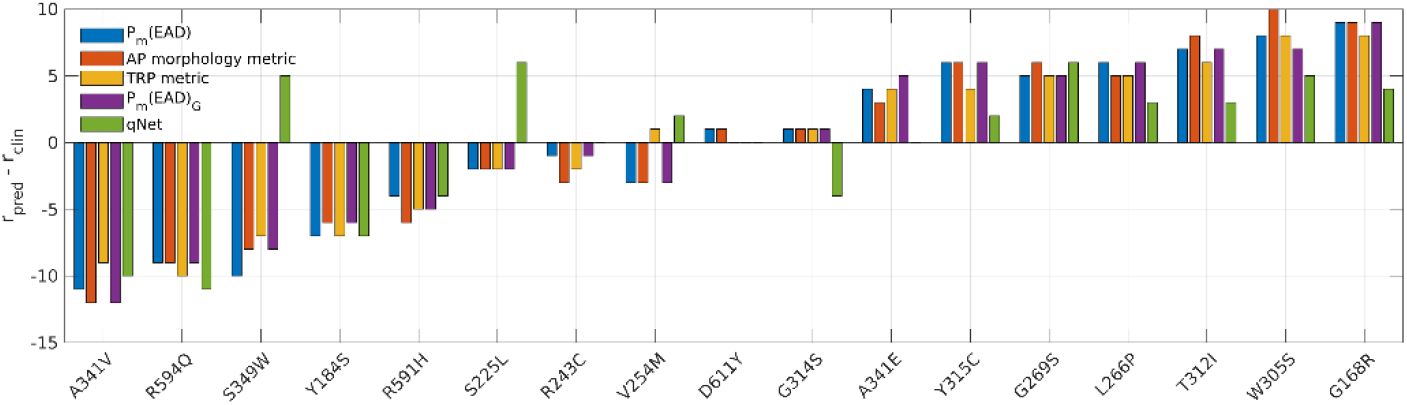
Ranking comparison across 5 metric – model pairs. Similar figure as Fig 3 except that the r_pred_ from P_m_(EAD)_G_ and qNet are included.

**S1 Table.**
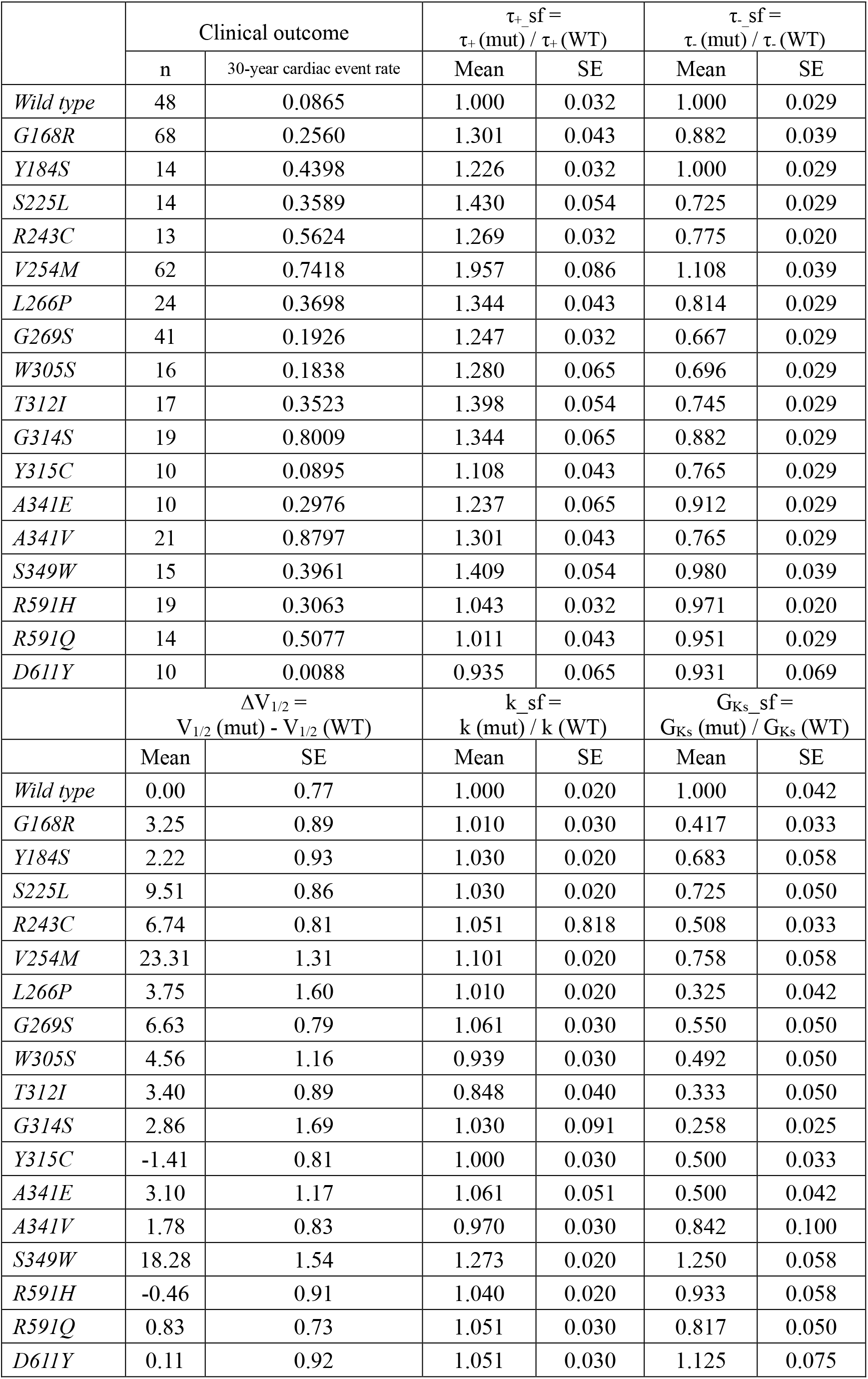
Long QT syndrome type 1 clinical data and cellular experimental data obtained from Jons et al., n: number of patients.

**S2 Table.** Five approaches prediction and ranking on 17 mutations

|  | Clinical data |  | P<sub>m</sub>(EAD) |  | P<sub>m</sub>(EAD)<sub>G</sub> |  |
| --- | --- | --- | --- | --- | --- | --- |
|  | risk | ranking | metric | ranking | metric | ranking |
| *D611Y* | 0.009 | 1 | 0.025 | 2 | 0.072 | 1 |
| *Y315C* | 0.090 | 2 | 0.260 | 8 | 0.573 | 8 |
| *W305S* | 0.184 | 3 | 0.404 | 11 | 0.625 | 10 |
| *G269S* | 0.193 | 4 | 0.347 | 9 | 0.604 | 9 |
| *G168R* | 0.256 | 5 | 0.551 | 14 | 0.819 | 14 |
| *A341E* | 0.298 | 6 | 0.383 | 10 | 0.651 | 11 |
| *R591H* | 0.306 | 7 | 0.033 | 3 | 0.120 | 2 |
| *T312I* | 0.352 | 8 | 0.664 | 15 | 0.850 | 15 |
| *S225L* | 0.359 | 9 | 0.199 | 7 | 0.488 | 7 |
| *L266P* | 0.370 | 10 | 0.699 | 16 | 0.873 | 16 |
| *S349W* | 0.396 | 11 | 0.024 | 1 | 0.143 | 3 |
| *Y184S* | 0.440 | 12 | 0.159 | 5 | 0.404 | 6 |
| *R594Q* | 0.508 | 13 | 0.051 | 4 | 0.180 | 4 |
| *R243C* | 0.562 | 14 | 0.548 | 13 | 0.806 | 13 |
| *V254M* | 0.742 | 15 | 0.436 | 12 | 0.759 | 12 |
| *G314S* | 0.801 | 16 | 0.900 | 17 | 0.962 | 17 |
| *A341V* | 0.880 | 17 | 0.172 | 6 | 0.304 | 5 |
|  | AP morphology |  | TRP |  | Qnet |  |
|  | metric | ranking | metric | ranking | metric | ranking |
| *D611Y* | 0.006 | 2 | -2.76 | 1 | 75.162 | 1 |
| *Y315C* | 2.313 | 8 | 19.66 | 6 | 74.460 | 4 |
| *W305S* | 3.731 | 13 | 29.41 | 11 | 73.403 | 8 |
| *G269S* | 2.883 | 10 | 26.34 | 9 | 73.212 | 10 |
| *G168R* | 3.990 | 14 | 33.43 | 13 | 73.321 | 9 |
| *A341E* | 2.814 | 9 | 27.93 | 10 | 73.508 | 6 |
| *R591H* | 0.000 | 1 | 6.33 | 2 | 74.631 | 3 |
| *T312I* | 5.666 | 16 | 37.62 | 14 | 73.190 | 11 |
| *S225L* | 1.893 | 7 | 22.05 | 7 | 72.401 | 15 |
| *L266P* | 5.289 | 15 | 38.36 | 15 | 73.148 | 13 |
| *S349W* | 0.024 | 3 | 15.5 | 4 | 71.428 | 16 |
| *Y184S* | 1.010 | 6 | 17.55 | 5 | 73.638 | 5 |
| *R594Q* | 0.133 | 4 | 12.44 | 3 | 74.688 | 2 |
| *R243C* | 3.161 | 11 | 30.55 | 12 | 73.119 | 14 |
| *V254M* | 3.620 | 12 | 43.92 | 16 | 70.040 | 17 |
| *G314S* | 5.811 | 17 | 46.89 | 17 | 73.183 | 12 |
| *A341V* | 0.308 | 5 | 24.56 | 8 | 73.490 | 7 |

**S3 Table.** Parameters chosen for P(EAD) prediction.

| parameters | Lower<br>bound | Upper<br>bound |
| --- | --- | --- |
| $\tau_+ \text{ sf}$ | 0.2903 | 2.8510 |
| $\tau_- \text{ sf}$ | 0.2051 | 1.6577 |
| $\Delta V_{1/2}$ | -11.9267 | 36.4100 |
| $k \text{ sf}$ | 0 | 10.8687 |
| $G_{K_S} \text{ sf}$ | 0 | 1.9044 |

