## Supplements for "Estimating the Probability of Early Afterdepolarization and Predicting Arrhythmic Risk associated with Long QT Syndrome Type 1 Mutations"

**Supplementary Methods**

**Authors:** Qingchu Jin^1^, Joseph L. Greenstein^1^, Raimond L. Winslow, PhD^1^

**Affiliations:**

^1^Department of Biomedical Engineering and Institute for Computational Medicine, Johns Hopkins University, Baltimore, MD, United States of America

### **Logistic regression model**

#### **Statistical modeling**

We described the detailed workflow of this statistical modeling approach previously [1]. In brief, we performed a two-stage random sampling strategy in the region of interest to obtain adequate sample parameter sets and then use these samples to fit the LRM (Eq (2)). In the first sampling stage, we uniformly randomly sampled 100 parameter sets. For each parameter set, 110 realizations with different random seeds are performed from which a value of P(EAD) for the parameter set can be calculated. Given the parameter sets and their corresponding P(EAD) values, a logistic equation mapping these parameters to P(EAD) is fitted. We used this equation to obtain a coarse estimation of the boundaries of the transition domain within the region of interest. Then, we uniformly randomly sampled an additional 100 parameter sets within the estimated transition domain, denoted as the second sampling stage. Similarly, 110 realizations are performed for each parameter set to get the corresponding P(EAD). Finally, the LRM is fitted by all parameter sets sampled from both stages. To improve the accuracy of the LRM, quadratic features derived from parameters are included in the LRM fitting process. The consistent Akaike information criterion (CAIC) is used to select quadratic features. To be specific, we tested all possible combinations of quadratic features and used the combination which yielded the highest CAIC on the two-stage parameter set samples as the LRM quadratic features.

#### **Region of interest of model parameters**

When building the LRM for predicting probability of occurrence of EADs in the setting of LQTS1, 5 I_Ks­_-related parameters were selected to be varied (S3 Table). These parameters were varied over an experimentally determined range of values (± 2 standard deviations) based on the work of Jons et al.[2] which we refer to here as the region of interest. Those 5 parameters are activation time constant scaling factor (τ_+__sf), deactivation time constant scaling factor (τ_-__sf), shift in the half maximal activation voltage (∆V_1/2­_), scaling factor of voltage-dependent activation curve slope (k_sf), and maximal conductance scaling factor (G_Ks__sf).

The region of interest can be separated into three mutually exclusive subspaces in terms of the arrhythmia event probability: lower plateau domain, upper plateau domain and transition domain. The lower plateau domain is the space where parameter sets simulated in the myocyte model yield P(EAD) estimated from myocyte model realizations (P(EAD)_MM_) that is strictly equal to 0 in this simulation (i.e., no EADs exhibited in the 110 realizations). In contrast, upper plateau domain is the space where P(EAD)_MM_ is strictly equal to 1 in this simulation (i.e., all 110 realizations exhibit EADs). The transition domain is the space where parameter sets simulated in the myocyte model yield P(EAD)_MM_ strictly > 0 and < 1.

#### **LRM Validation Metrics**

Linear regression was performed between myocyte model-generated (actual) values of P(EAD) and LRM predicted values of P(EAD). The R-squared (R^2^) from the linear regression was calculated. Absolute error between those was also calculated as

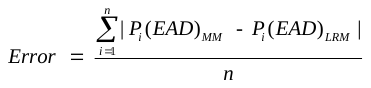
 (S1)

,where *n* is the number of parameter sets, and P_i_(EAD)_MM_ and P_i_(EAD)_LRM_ are event (i.e. EAD) probabilities associated with the *i*th parameter set obtained from the myocyte model and LRM, respectively.

#### **Uncertainty Analyses**

The LRM derived in this study is a function that maps parameters to P(event). If experimental measurements of these parameters are variable, then this variation can be modeled by assuming these measurements are drawn from an underlying probability distribution. In this case, the event probability produced by the LRM is also a random variable (RV). We wish to characterize this uncertainty by computing the distribution of the estimated event probability. To do this, when the mean and variance of parameter estimates are available from experimental data, we assume parameters are RVs drawn from a normal distribution with that mean and variance. To ensure the values are limited to be within a biophysically meaningful range for each parameter, distributions were truncated to exclude values outside this range. We randomly generate (10^6^) sample vectors for a specific parameter set based on the underlying parameter uncertainty distributions and use the LRM to predict P(EAD) for all sample vectors. Following this approach, the P(EAD) distribution can be estimated.

### **Implementation details of model-based risk metrics**

#### **P_m­_(EAD)_G_ metric.**

To test the generalizability of the mean P(EAD) (P_m­_(EAD)) metric, we built a new LRM using the Greenstein stochastic myocyte model [3] to calculate P_m_(EAD). We denote P_m_(EAD) obtained using the Greenstein model as P_m_(EAD)_G_.

To incorporate I_Ks_-related parameters, we incorporated the Jons et al. [2] I_Ks_ model into the Greenstein model. To be consistent with the P_m_(EAD) computed from the Walker model, we implemented the β-adrenergic stimulation protocol described in Greenstein et al. [4]. Several modifications are made to the β-adrenergic stimulation protocol to increase the proarrhythmic effect. 1) The 3.3-fold increase on SERCA is reduced to 2-fold. 2) L-type Ca^2+^ channel opening rate scaling factor is changed from 0.2 to 0.7. 3) The β-adrenergic stimulation protocol for I_Ks_ upregulation changed from the 2-fold increase of G_Ks_ to the followings: reducing half activation voltage from 9.4mV to 44.4mV combined with a 1.4-fold increase of G_Ks_.

We used the same region of interest described in S3 Table to build the LRM for the Greenstein model. For simplicity, we only performed the first stage sampling described in statistical modeling section and Jin et al. [1]. 100 parameter sets are randomly sampled in the region of interest and ~250 realizations are performed for each parameter set. The simulation protocol described in the Methods is used for each realization except that each realization lasts for 1 second. The initial state of all realizations is the diastolic state following the 20^th^ beat under β-adrenergic stimulation with the wild type I_Ks_. To detect the EAD, we estimated the local minima and local maxima during the plateau phase of the action potential (AP). If the difference between the greatest local maxima and the smallest local minima is larger than 5mV, then we consider the AP to have triggered an EAD. For simplicity, we used the same features described in Table 1 to build the LRM. The LRM has a good performance with an error of 0.017 ± 0.024 P(EAD) on these 100 parameter sets. Then, with the LRM based on the Greenstein model, we calculated the P_m_(EAD)_G_ for the 17 I_Ks_ mutations described in S1 Table.

#### **AP morphology metric.**

Kernik et al. leveraged a population model of induced pluripotent stem cell derived cardiomyocytes (iPSC-CMs) to develop an approach for LQTS1 arrhythmia risk prediction [5]. In this approach, for each mutation, the wild-type I_Ks_ is optimized to fit to I_Ks_-related parameters. The percentage of models in the population model for each mutation that satisfy all of the following 3 conditions is taken as the risk metric: 1) exhibiting greater than 4% increase of the APD_90_, 2) exhibiting at least 4% increase of beat-to-beat variation, and 3) exhibiting at least 4% increase of APD triangulation (APD_90_-APD_30_) [6]. The larger the percentage, the greater risk the mutation is considered to be. We denote this metric as AP morphology metric.

The simulation process is the same as the description in Kernik et al. [6] with several additional constraints. 1) The steady state was defined by <1% change in minimum ion concentrations between the first and last beat of a 50s simulation run. If the model is still unable to achieve <1% change at 700^th^ beat, the state at the 700^th^ beat is considered as the initial state. 2) During the calculation of APD_90_, the start of the AP is defined as the time point when the
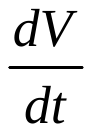
 first reaches 90% of the max
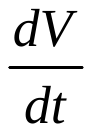
 of the entire AP profile. 3) In the AP morphology analysis, only APs satisfying the inclusion criteria are studied, where the inclusion criteria require that AP amplitude is over 70mV and max
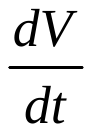
 is larger than 1mV/ms. This ensures that cells were beating and fully repolarizing.

#### **qNet.**

qNet is an extensively studied metric for drug proarrhythmic potential prediction, developed in the Comprehensive in Vitro Proarrhythmia Assay group guided by the Food and Drug Administration [7]. qNet calculates the integral of 6 currents over 1 AP: L-type Ca^2+^ current, late Na^+^ current, fast K^+^ current, I_Ks_, inward rectifier K^+^ current, and transient outward K^+^ current. The modified O’hara Rudy model [8] was used as the platform to calculate qNet. The risk of mutation is increases as qNet value decreases.

To incorporate I_Ks_-related mutation parameters, we incorporated the Jons et al. [2] I_Ks_ model into their modified O’hara Rudy model using a value of GKs for wild-type matching that originally used in the modified O’hara Rudy model in order to maintain the correct AP morphology. For each mutation, an AP train 1000 beats (cycle length = 2000ms) is performed. The beat having the largest
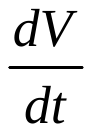
 out of the last 250 beats is used to calculate qNet.

All Models and results are available in Github (<https://github.com/JHU-Winslow-Lab/EAD-paper-code.git>).
